# Power asymmetry destabilizes reciprocal cooperation in social dilemmas

**DOI:** 10.1101/2024.09.02.610740

**Authors:** Marco Colnaghi, Fernando P. Santos, Paul A. M. Van Lange, Daniel Balliet

**Author notes:** Corresponding authors. (MC), (DB). These authors contributed equally to this work (Fernando P. Santos, Paul A. M. Van Lange). **Author contributions:** Conceptualization: MC, DB, FPS, PAMVL; Methodology: MC, DB, FPS, PAMVL; Software: MC; Formal Analysis: MC; Writing—original draft: MC; Writing— review & editing: MC, DB, FPS, PAMVL; Visualization: MC; Funding acquisition: DB. **Competing interests:** All other authors declare they have no competing interests. **Code availability:** The code used to generate the results of this study has been deposited and is available on Zenodo (https://zenodo.org/doi/10.5281/zenodo.10687584).

## Abstract

Direct reciprocity has been long identified as a mechanism to enhance cooperation in social dilemmas. While most research on reciprocal cooperation has focused on symmetrical interactions, real world interactions often involve differences in power. Verbal theories have either claimed that power differences enhance or destabilize cooperation, indicating the need for a comprehensive theoretical model of how power asymmetries affect direct reciprocity. Here, we investigate the relationship between power and cooperation in two frequently studied social dilemmas, the prisoner’s dilemma (PD) and the snowdrift game (SD). Combining evolutionary game theory and agent-based models, we demonstrate that power asymmetries are detrimental to the evolution of cooperation. Strategies that are contingent on power within an interaction provide a selective advantage in the iterated SD, but not in the iterated PD. In both games, the rate of cooperation declines as power asymmetry increases, indicating that a more egalitarian distribution of the benefits of cooperation is the prerequisite for reciprocal cooperation to evolve and be maintained.

## Background

Social dilemmas, where individuals can increase their payoffs at the expense of collective welfare, present an important challenge to cooperation (1–4). When social interactions between individuals are repeated, direct reciprocity can provide a strong incentive towards cooperative behaviour (5–10). If the continuation probability (i.e., the likelihood of interacting again with the same individual) exceeds a critical threshold, reciprocal strategies such as Tit-For-Tat (TFT: cooperate on the first encounter, then cooperate only if one’s social partner cooperated on the previous round, and defect otherwise) can outcompete defection, paving the way for cooperation to spread in a population of free-riders (7,11).

Most research on direct reciprocity in dyadic interactions has focused on symmetric games (8–13). Yet, many real-world interactions are characterized by some form of power asymmetry, where two (or more) persons differ in how strongly they can influence each another’s outcomes (14–16). Humans can readily infer and respond to power asymmetries across social interactions (17–21). Power differences are widespread in the animal kingdom as well (22–24), and the role of power asymmetry in animal contests has been investigated by several theoretical studies (25–27). However, it is only in recent years that researchers have begun to study the impact of power asymmetries on direct reciprocity (28–30). Here, we offer a theoretical framework for understanding why, and under what conditions, power may or may not undermine the necessary conditions for cooperation.

### Power asymmetry and cooperation

Power can be broadly defined as asymmetric control over another person’s outcomes (20,31,32). Experimental and theoretical studies on power asymmetry typically focus on one (or a few) of a number of factors that can lead to asymmetric control, such as differences in payoffs (33–36), endowments (29), effectiveness of punishment (37,38), or the ability to choose one’s social partner (39). Empirical studies that consider differences in payoffs indicate that asymmetry can pose a barrier to the emergence of cooperation in social dilemmas (33–35,39). Previous theoretical research also suggests that differences in power might destabilize cooperation in social dilemmas (28) and indicates that inequality can undermine cooperation in public goods games (29). At the same time, the effects of power differences are not unequivocal or universal; power asymmetries do not always undermine (or promote) cooperation (37,40) and power can yield different effects on cooperation in different countries (41). In fact, power hierarchies have even been considered functionally adaptive by promoting cooperation within a group (42,43). Here, we operationalize power asymmetry in a social interaction as the ability to provide higher or lower benefits to a partner in a simultaneous donation game (20,44,45), and in doing so we aim to clarify the relationship between power asymmetry and cooperation.

### Inference of power asymmetry

In a social ecology with power differences across interactions, there could exist adaptive benefits to inferring power differences and conditioning behavioural strategies of cooperation on power. According to functional interdependence theory (FIT), power is one of several dimensions of interdependence that humans can infer to decide when and with whom to cooperate (46,47). FIT posits that the ability to infer and respond to power asymmetries (and other dimensions of interdependence, such as correspondence or conflict of interests) is an adaptation that provided a selective advantage in our ancestral environment (46). Indeed, humans can readily detect, and respond to, differences in power (17–21), and people employ a wide range of visual and auditory cues to infer differences in power (48–50). Children as young as five can accurately use nonverbal behaviour to discriminate between high- and low-power individuals (51), and perceived power differences can, in turn, induce major changes in one’s affective and cognitive state (21,52,53). In the context of symmetric interactions (i.e., without any differences in power), it has been shown that heterogeneous social environments can select for the ability to infer interdependence (54). In the more general case of asymmetric interactions, however, little is known about the features of the social environments that promote adaptations to infer power differences.

In the present study, we analyse the interplay between power asymmetry (in the form of unequal resource allocation) and reciprocal cooperation in iterated 2-person games. We aim to clarify how power affects reciprocal cooperation, and under what conditions can the ability to infer power differences provide a selective advantage. We start by studying reciprocal cooperation in the iterated, asymmetric version of the simultaneous donation game, a specific form of the Prisoner’s Dilemma (PD). We derive a simple formula, which expresses the minimum continuation probability necessary to stabilize cooperation as a function of power asymmetry. We then consider strategies that are conditional on power differences within interactions, and evaluate whether the ability to adapt one’s behaviour in response to power asymmetry is evolutionarily stable and provides a selective advantage in the iterated PD. We subsequently shift our attention to the asymmetric version of the iterated Snowdrift game (SD; also referred to as Chicken game), another frequently studied social dilemma (9,55). While the dynamics of the iterated PD become similar to a Stag Hunt (SH) (56) if the continuation probability is high enough, the dynamics of the iterated SD between reciprocal cooperators resemble a maximizing difference/harmony game (see Methods). Therefore, focusing on the iterated SD allows us to extend our analysis to all four “archetypal” games that people most frequently use to describe social interactions (57) and are most frequently studied in the literature on social dilemmas (54,58,59). Additionally, in the SD, individuals can increase their payoffs by doing the opposite of what their social partner does; in line with previous studies, we refer to this behaviour as “anti-coordination” (see, for example, ref.(60)).Therefore, the SD allows us to study the evolution of cooperation in a situation where differences in power can help solve collective action problems (61–63).

## Results

### A simple rule for the evolution of reciprocal cooperation in an asymmetric PD

First, consider the simple case of two individuals, who differ in their level of power, interacting with each other through the asymmetric simultaneous donation game. In the symmetric version of this game, each individual can incur a cost to provide a benefit to their social partner (64). We consider an asymmetric version of this social dilemma, where high-power individuals confer a higher benefit, (1 + *α*)*b*, and low-power individuals a lower one, (1 ™ *α*)*b*, with 0 ≤. *α* < 1 By introducing and *α* assuming this distribution of payoffs, we guarantee that power asymmetries to not affect the overall benefits of cooperation. We assume the cost of cooperation to be the same for high- and low-power individuals. This leads to the following payoff matrix:

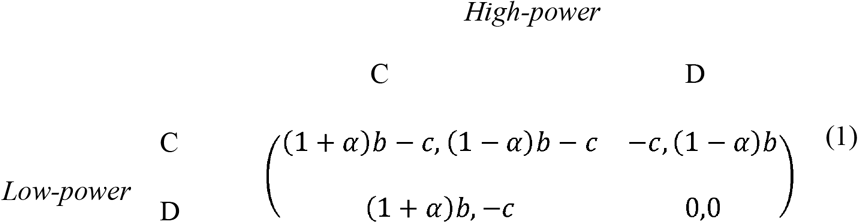

Using the framework of interdependence theory (46,47), it is possible to show that, for the parameters used throughout the study, the degree of asymmetric dependence (a method of assessing power asymmetry) increases monotonically with *α* (Methods; Figure S1). In other words, the greater the asymmetry (*α*), the more a low-power individual’s outcome is influenced by their partner’s choice to cooperate or not. For simplicity, throughout the paper we refer to *α* as power asymmetry. This definition of power is consistent with theoretical work that suggests that the ability to allocate greater rewards is an expression of power (20,44,45).

In order to study the impact of direct reciprocity, we assume that there is a continuation probability *W* of repeated interaction with the same individual, and individuals can either choose reciprocal cooperation (TFT) or always defect (AllD). We limit our analysis to these two strategies as we are interested in the necessary conditions that promote the evolution of reciprocal cooperation in a population of defectors; if TFT cannot displace AllD, no other strategy can (7). Moreover, when cooperation costs are high, TFT remains a key strategy to sustain cooperation even when stochastic and longer-memory strategies are considered (65). When only these two strategies are considered, it is possible to prove (see Methods) that TFT is an evolutionarily stable strategy (ESS) if, and only if, the continuation probability exceeds the threshold:

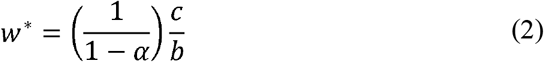

For *α =* 0, we retrieve the well-known condition for the evolution of cooperation in the symmetrical PD, *w*,> *c/b* (7). When *α >* 0 any increase in power asymmetry leads to a higher threshold continuation probability, making it harder for cooperation to evolve (Figure 1). As *α* approaches the value of 1, the minimum continuation probability necessary for cooperation to be stable diverges to infinity. This simple formula illustrates the negative impact that power asymmetry has on cooperation: even if *w* is high enough to sustain cooperation in the symmetric case, increasing power inequality will eventually lead to the breakdown of cooperation.

**Figure 1.**
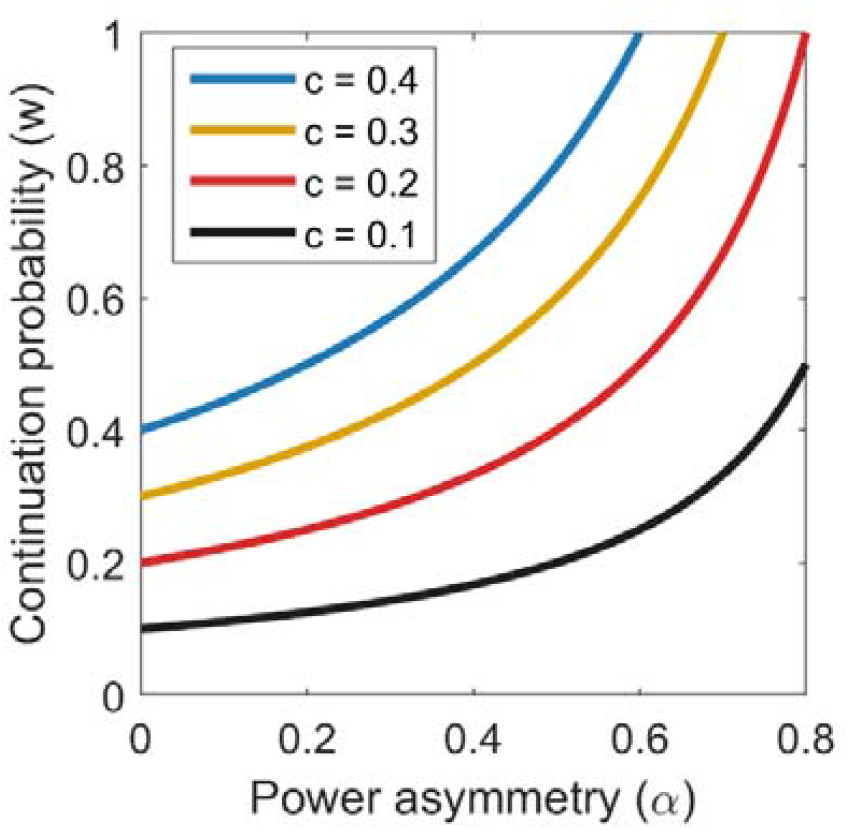
Threshold continuation probability. Minimum continuation probability *w* * that makes reciprocal cooperation (TFT) evolutionarily stable. Increasing power asymmetry (α) or the cost of cooperation (*c*)destabilizes cooperation. Other parameters: *b*=1.

### Evolutionary stability and fixation probability

Next, we consider the more complex case of a population where individuals have an equal probability of finding themselves in a position of high or low power. This assumption reflects the empirical observation that most variation in perceived power is due to changes in situations, rather than stable individual traits (21). Individuals can therefore find themselves in three types of possible interactions: symmetric, low-power (when a low-power individual interacts with a high-power one), and high-power (when a high-power individual interacts with a low-power one). To study whether the ability to infer differences in power can provide a selective advantage in the PD, we consider eight possible strategies: two “power-independent” strategies, where individuals always play TFT or AllD in every situation regardless of power asymmetries, and six strategies conditional on power, where individuals choose whether to play TFT or AllD depending on the specific type of interaction (symmetric or with a higher/lower-power individual). Each of the eight strategies can then be identified as a triplet (*X,Y,Z*) where an individual plays *X* in symmetric interactions, *Y* when in low power and interacting with a higher-power individual, and *Z* when in high power and interacting with lower-power individual. From simplicity, we use the shorthand “C” to refer to reciprocal cooperation (TFT), and “D” as a shorthand for AllD. For example, the strategy (*C,C,D*) plays TFT in symmetric interactions and when interacting with higher-power individuals, and AllD when interacting with individuals with a lower power status.

Using a standard evolutionary game-theoretical approach (66), we evaluate under what conditions each of these strategies is evolutionarily stable (Figure 2). In addition, we use a small-mutation approximation (67) to evaluate whether each of the strategies that are evolutionarily stable in an infinite population would evolve in a finite population subject to stochastic perturbations (Figure 3). The ESS analysis shows that, for a given level of power asymmetry, a cooperative power-independent strategy (*C,C,C*) is evolutionarily stable, provided that the continuation probability is high enough (Figure 2). As power asymmetry increases, the minimum threshold that makes reciprocal cooperation advantageous increases as well, making cooperation less stable. As a single cooperative mutant can never invade an infinite population of defectors, (*D, D, D*) is an ESS in the whole parameter space (Figure 2). In addition to these two power-independent strategies, two mixed strategies, (*D, C, C*,) and (*C,D,D*), are evolutionarily stable in two regions of the parameter space. However, a finite-population size analysis reveals that these two strategies almost never outperform power-independent ones (Figure 3). (*C,D,D*) outperforms other strategies only for a limited range of continuation probability *W* and in large populations (*N* = 500,1000; Fig. S2). (*D, C, C*) is never advantageous in a finite population. In principle, these strategies are stable once they spread to fixation in an infinite population; in practice, however, natural selection does not promote their spread (if not in a very narrow range of ecological conditions), making the fixation of power-independent strategies more likely. When repeated interactions are infrequent, natural selection favours defection in all situations (*D, D, D*) (Figure 3). As the continuation probability increases, so does the equilibrium frequency of (*C, C, C*), eventually outcompeting (*D, D, D*): once cooperation becomes advantageous, it is favorable to cooperate in all situations, regardless of whether the interaction is symmetric or asymmetric (Figure 3). To conclude, the ability to infer power and modify one’s behaviour accordingly is unlikely to provide a selective advantage in the iterated asymmetric Prisoner’s Dilemma.

**Figure 2.**
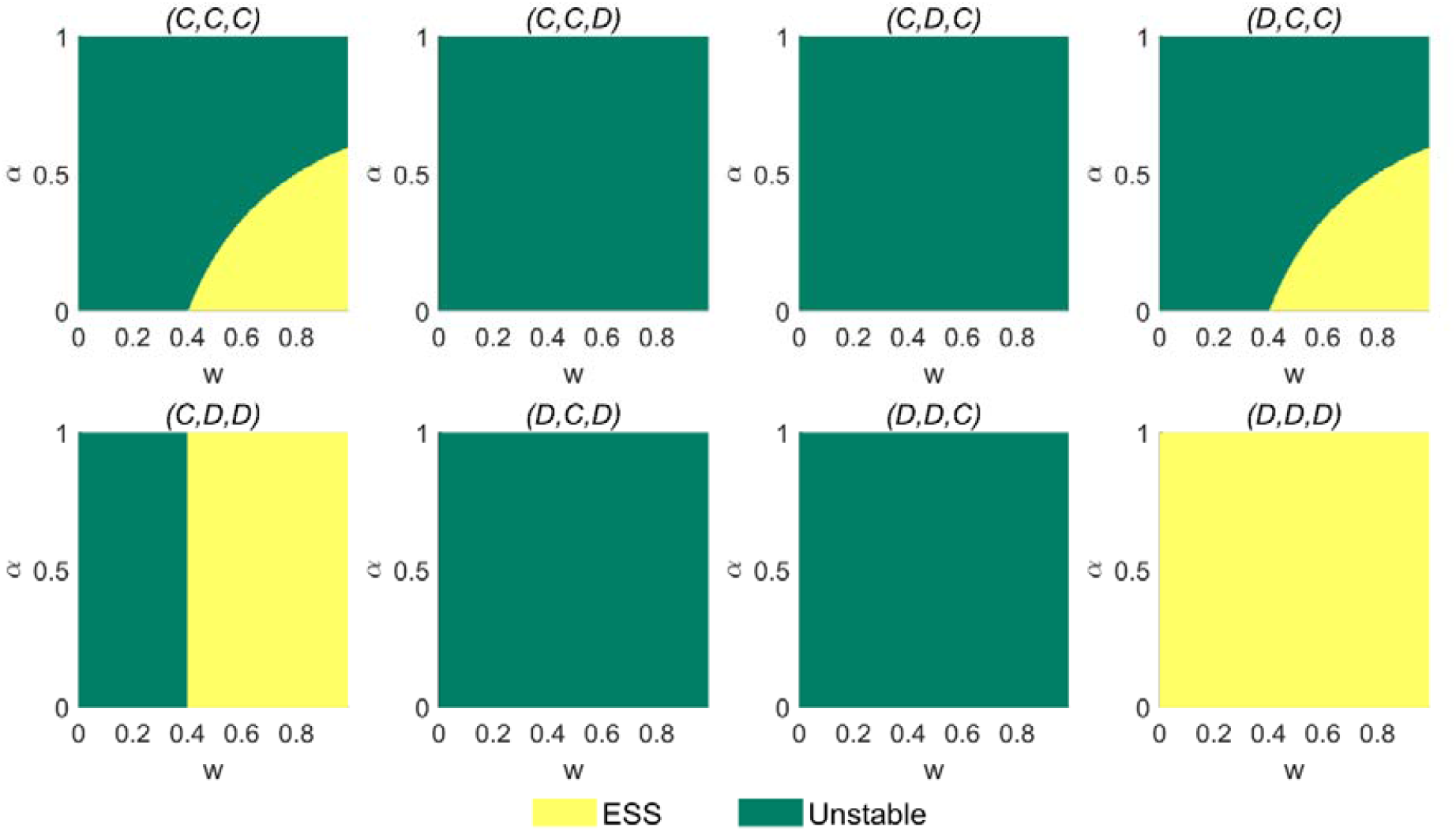
Evolutionarily stable strategies in the iterated PD. Evolutionarily stable strategies in the iterated Prisoner’s Dilemma for varying levels of power asymmetry (α) and continuation probability (*w*) The area of the parameter space where each strategy is an ESS is indicated in yellow. Other parameters: *b=*1,*c*=0.4.

**Figure 3.**
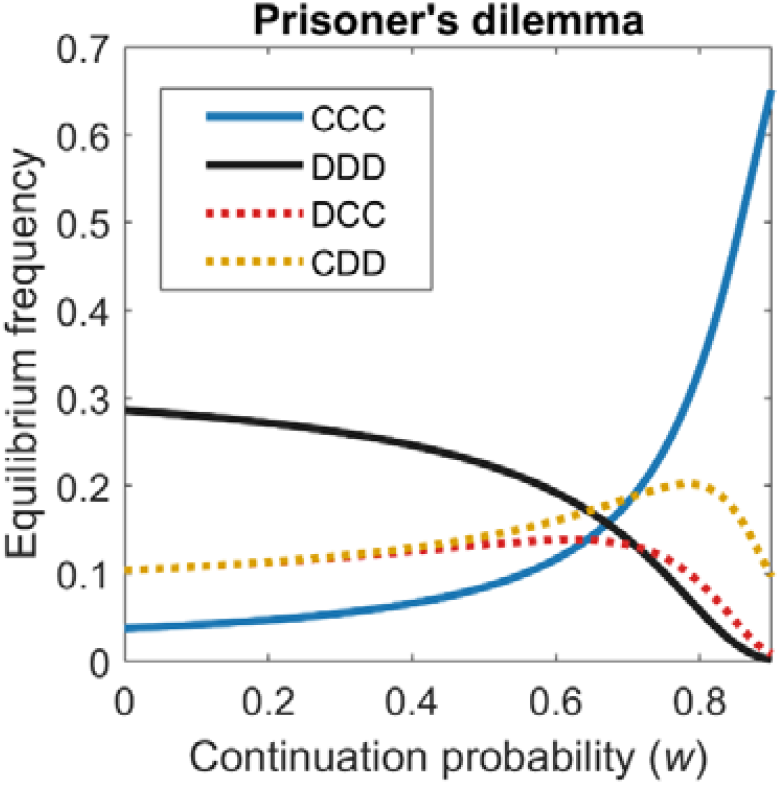
Equilibrium frequency of strategies in the iterated PD. Equilibrium frequency of various strategies in a finite population playing the iterated PD, under a small-mutation approximation, as a function of the continuation probability (*w*). Continuous lines indicate power-independent strategies; dotted lines indicate strategies that are conditional on power differences. Of the 8 strategies studied, only the four ESSs are shown. Other parameters: α=0.5, *b=*1,*c*=0.4,*N*=100, *β*=0.05

### Asymmetric Snowdrift game

The simultaneous donation game described above can be modified to assume the form of a Snowdrift (SD) game, a slightly more benign social dilemma where cooperators and defectors can stably coexist in a population (9,55). This game represents a situation where two individuals must work together to achieve a certain outcome, such as building a shelter or freeing a road from a snowdrift, which benefits them both. Each individual would be better off if their partner would bear the whole cost of this enterprise, but incurring the cost of cooperation alone is preferable to the situation where neither player contributes to the common good.

Let 2*b* be the total payoff obtained through mutual cooperation. Suppose that the total cost of the endeavor, 2*c* can be either split between the two individuals or borne by one individual alone. As in the previous case, the reward can be either divided symmetrically when two individuals of equal power meet, or asymmetrically, if there is a power difference between the two players. In the former case, both players receive the same reward. *b* In the latter, the high-power player shares with the other a greater proportion of the reward, (1+ *α*)*b* and takes a smaller proportion, (1 ™ *α*)*b* for themselves. This specific form of SD game allows us to compare the results directly with the asymmetric PD, as the resulting payoff matrix has the same reward for mutual cooperation:

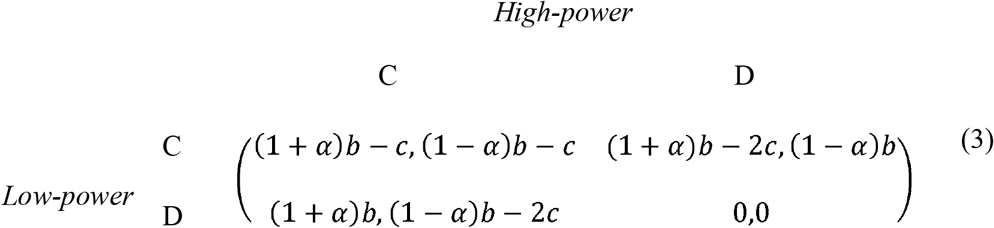

Again, we assume a continuation probability *w* of repeated interaction with the same partner, and consider two different behaviors: reciprocal cooperation (TFT) or always defect (AllD). In the simple case of a high-power individual interacting with a low-power one, we retrieve the same simple rule expressed by Equation (2). In the more complex case of a population where individuals can find themselves in a high-or low-power position with equal probability, we consider the two power-independent and the six strategies conditional on power introduced in the previous section (see Methods). Again, we evaluate which strategies are ESSs, and apply a small-mutation approximation to analyse which of the ESSs are favoured by natural selection in a finite population.

A standard evolutionary game-theoretical analysis (66) reveals that (as in the case of the PD) a fixed strategy of reciprocal cooperation (*C, C, C*) becomes evolutionarily stable once the continuation probability is high enough and power asymmetry is not too extreme (Figure 4). As power asymmetry increases, (*C, C, C*) becomes unstable and is replaced by a strategy conditional on power differences, (*C, C, C*), which plays TFT in symmetric interactions and when interacting with higher-power individuals, and defects when interacting with lower-power individuals (Figure 4). Performing a small-mutation, finite population size analysis, we find that this strategy outcompetes both power-independent strategies, (*C, C, C*) and (*D, D, D*), when the continuation probability is low (Figure 5). As *w* increases, (*C, C, C*) becomes more favorable, and it eventually outcompetes (*C, C, D*) (Figure 5). These conclusions are not affected by population size (Figure S3). Thus, if the continuation probability does not exceed this threshold, the ability to infer power and modify one’s behavior accordingly provides a selective advantage in the iterated asymmetric SD.

**Figure 4.**
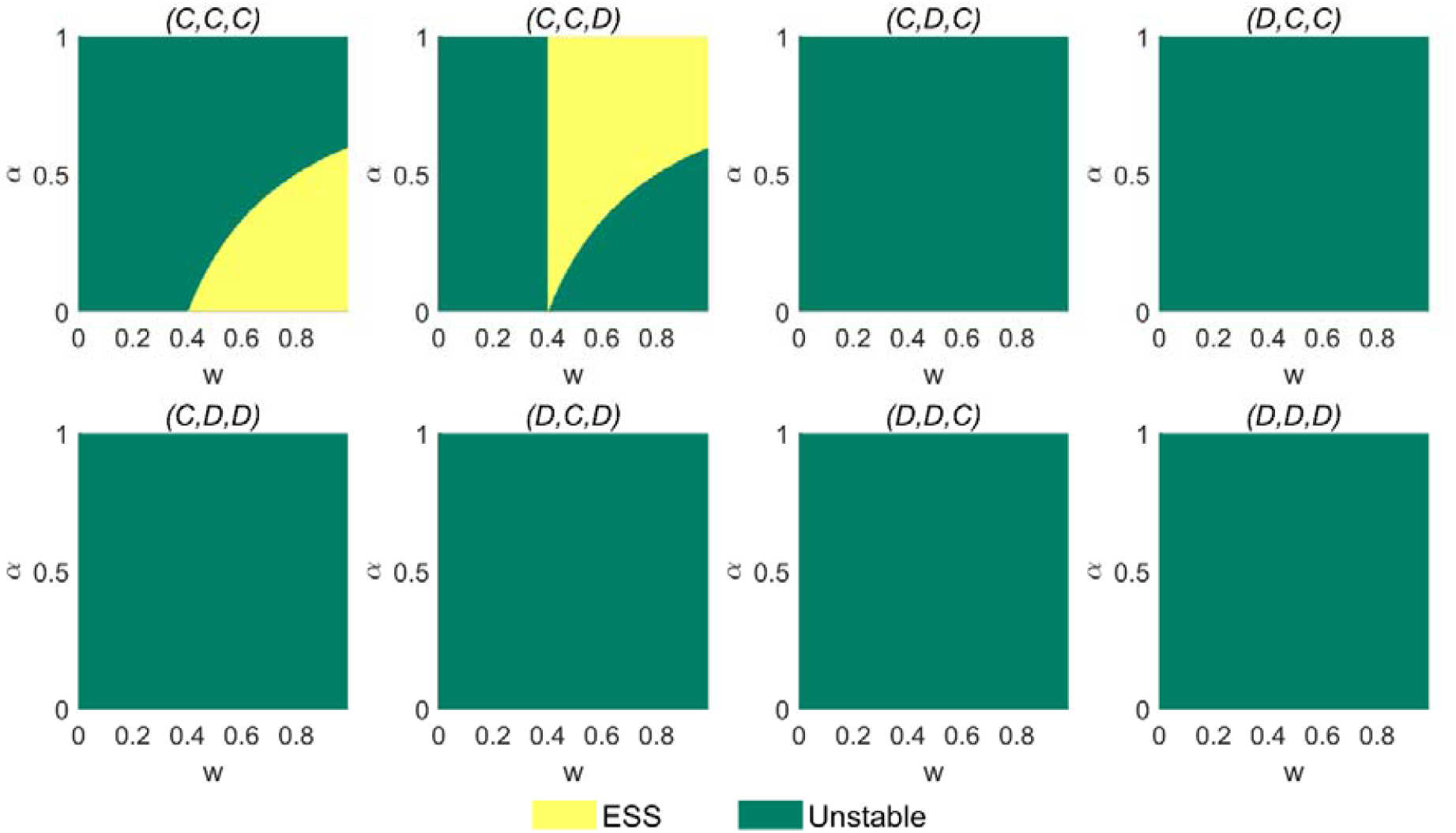
Evolutionarily stable strategies in the iterated SD. Evolutionarily stable strategies in the iterated Snowdrift for varying levels of power asymmetry (α) and continuation probability. (*w*)The area of the parameter space where each strategy is an ESS is indicated in yellow. Other parameters :*b=*1,*c*=0.4.

**Figure 5.**
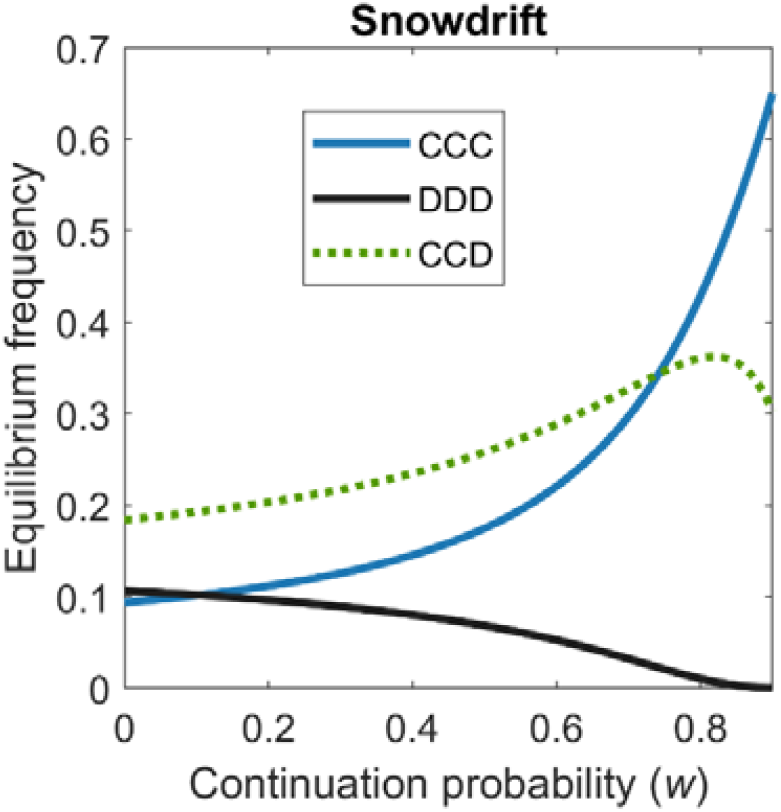
Equilibrium frequency of strategies in the iterated SD. Equilibrium frequency of various strategies in a finite population playing the iterated SD, under a small-mutation approximation, as a function of the continuation probability (*w*). Continuous lines indicate power-independent strategies; the dotted line indicates (*C,C,D*), a strategy that is conditional on power differences. Of the 8 strategies studied, only the two ESSs and AllD are shown. Other parameters: α=0.5, *b=*1,*c*=0.4 *N*=100, *β*=0.05.

### Changes in cooperation frequency

Finally, we consider how the introduction of strategies conditional on power changes the average level of cooperation in a finite population under a small-mutation approximation, calculated as the equilibrium frequency of a strategy times the fraction of interactions where that strategy will play TFT (Figure 6). These results confirm our claim that cooperation declines with power asymmetry (Figure 6). This effect becomes more pronounced when the continuation probability is higher, and the decline in average cooperation increases with Thus, higher levels of power asymmetry cause a decline in the frequency of cooperative interactions.

**Figure 6.**
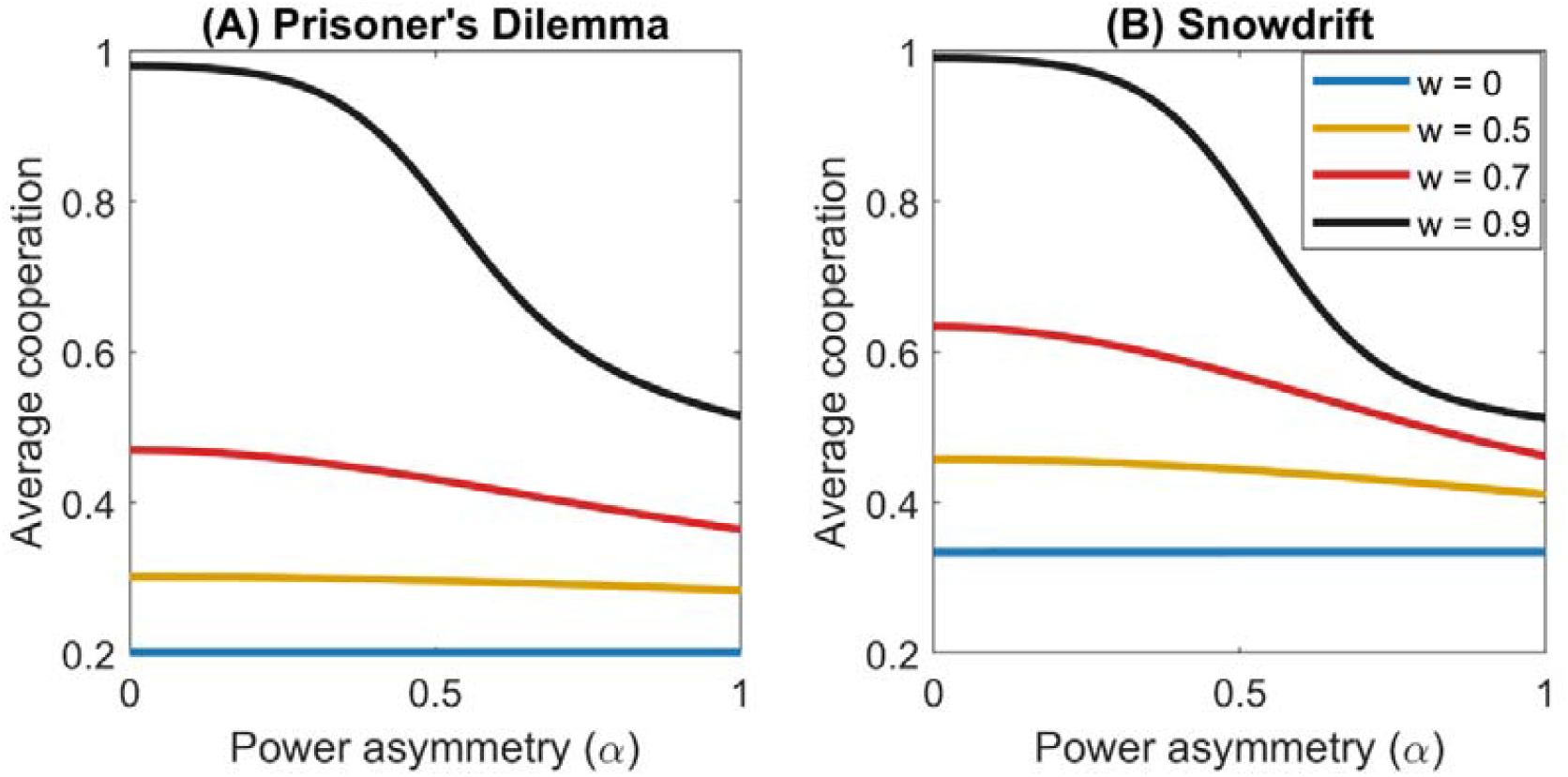
Average frequency of cooperation. Average frequency of cooperation in a finite population under a small-mutation approximation, as a function of power asymmetry (α) and the continuation probability (*w*), in the iterated PD (A) and SD (B). Other parameters: α=0.5, *b=*1,*c*=0.4 *N*=50, *β*=0.01.

## Discussion

In the symmetric prisoner’s dilemma (PD), direct reciprocity is evolutionarily stable if, and only if, the continuation probability (i.e., the likelihood of repeated encounters with the same individual) exceeds the cost-to-benefit ratio (7). We show that power asymmetry, modeled as the ability to allocate smaller or greater rewards in a simultaneous donation game, destabilizes reciprocal cooperation, raising the minimum continuation probability needed for cooperation to evolve (Equation 2, Figure 1). An intuitive explanation for this phenomenon is that the cost-to-benefit ratio of cooperation for high-power individuals (who can allocate greater rewards, and receive smaller ones) increases with power asymmetry: if the level of asymmetry exceeds a critical threshold, the reward for mutual cooperation is so small that exploiting one’s social partner once becomes more advantageous than sustained reciprocal cooperation. Therefore, it is in the interests of high-power individuals to defect, even when repeated interactions afford the possibility of reciprocal cooperation.

The ability to infer power differences across different interactions (symmetric, low-power, and high-power) does not provide any selective advantages in the PD. In this social dilemma, power-independent strategies always outcompete strategies that are contingent on power asymmetries, except for a very narrow range of continuation probability (Figure 3). By contrast, in the SD, a strategy that cooperates in symmetric and low-power interactions and defects in high-power interactions, (*C, C, D*), is evolutionarily stable in infinite populations (Figure 4) and favored by natural selection in finite populations (Figure 5) for low to intermediate values of the continuation probability. Among the strategies that are conditional on power, (*C, C, D*) is the one that achieves the highest payoffs: when (*C, C, D*) interacts with itself in asymmetric interactions, the low-power individual cooperates, and the high-power individual therefore receives the highest payoff available to them. This strategy can outcompete (*C, C, C*) by exploiting it in low-power interactions, and (*D, D, D*) by cooperating with itself in symmetric interactions. Perhaps less obviously, (*C, C, D*) can also outcompete other strategies conditional on power differences, such as (*C, D, C*). For example, a rare (*C, C, D*) mutant can invade a (*C,D,C*) population, but not vice versa, as the payoff of a low-power individual cooperating with a high-power individual is higher than that of a high-power individual cooperating with a low-power one.

This result suggests that the ability to infer differences in power provides a selective advantage in environments characterized by social dilemmas where anti-coordination is beneficial (i.e., when individuals can increase their payoffs by doing the opposite of what their partner does), such as the SD. In these games, the total payoffs are higher if individuals make opposite choices, especially because unilateral defection yields the best possible outcome whereas mutual defection yields the worst possible outcome (46,47). By contrast, when the continuation probability is high, the iterated PD takes the form of a Stag Hunt (56); in this type of game, the total payoffs are higher if individuals make matching choices, especially because mutual cooperation yields the best possible outcome whereas unilateral cooperation yields the worst possible outcome.

It has been suggested that the emergence of power asymmetries, such as in the form of leadership and followership, provided a selective advantage in ancestral human societies, as it helped to solve collective action problems (61–63). Repeated social interactions, in a context where a mixed group of leaders and followers performs better than a uniform group, give rise to dynamics typical of the SD: if everyone strives to be a leader, an individual can increase their own (and collective) payoffs by being a follower; in a population of followers, becoming a leader will also lead to higher payoffs. Our model indicates that, under these conditions, the ability to infer differences in power helps maximize joint payoffs by facilitating anti-coordination, and the evolution of decision rules based on power can therefore be promoted by natural selection. We do not observe this in the iterated PD, where individuals can increase their payoffs by matching their partner’s choice. In these games, strategies that discriminate based on power perform poorly against themselves, because they involve doing the opposite of what one’s partner does when interactions are asymmetric.

Power asymmetries are often addressed by cues that are shared between people or other animals. Social status is an example of cue that can be used to coordinate in social interactions. Prior research shows that people tend to defer to the preferences of higher-status individuals, even when status is arbitrarily assigned (68–70). This has been found to be particularly advantageous in asymmetric games, such as the battle of the sexes, where coordination would otherwise be difficult to achieve (70). While social status and power are distinct constructs, they are closely associated, and cues of status can be used to infer power (19,20).

Empirical studies on asymmetric games support the claim that asymmetry destabilizes cooperation (33–36,39). In asymmetric PDs, cooperation rates are significantly lower than in symmetric ones (34,35). Moreover, information about the payoff matrix reduces cooperation rate in asymmetric games (36), while increasing it in symmetric ones (36,71). Experimental studies also indicate that the temptation to defect is stronger for the players who have less to gain from cooperation (corresponding to the high-power players in our model) (34–36). By contrast, in repeated asymmetric games, players who receive higher payoffs (corresponding to low-power individuals in our model) are more likely to initiate cooperation and less likely to defect in response to their partner’s defection (34). These results indicate that, in agreement with our theoretical predictions, individuals who have less to gain from cooperation aim to minimize payoff disparity through frequent defection, thereby undermining the emergence of stable cooperation. While this behaviour is a suboptimal strategy in the PD (Figure 3), it can provide a selective advantage in the asymmetric SD (Figures 4 and 5).

Our theoretical results suggest further experimental studies to investigate whether the tendency to defect when in the high-power state is stronger in asymmetric SD than PD games, which would support the hypothesis that this behaviour confers a selective advantage in games where anti-coordination is on average beneficial. While the empirical evidence discussed above indicates that payoff differences hinder cooperation, there is no universal consensus on the effects of power, and different sources of power asymmetry may yield distinct outcomes (37,40,41). For example, differences in the effectiveness of punishment promote cooperation in a modified version of the PD, where players can contribute a variable investment to the common good (37), but not in a standard PD, where players only have a binary choice to cooperate or defect (38). The asymmetric ability to punish is also more effective in promoting cooperation when used to encourage joint cooperation, rather than to exploit one’s partner, in PD experiments (41). Different forms of asymmetry can also act synergistically with one another, as shown by a recent theoretical analysis of asymmetric public goods games (29). This study shows that extreme power asymmetry, in the form of an unequal distributions of endowments, prevents the emergence of reciprocal cooperation; but when the rewards of cooperation are also asymmetric, unequal endowments actually promote, rather than hinder, cooperative behaviour (29). Taken together, these studies suggest future theoretical and empirical work to further examine how different bases of power impact cooperation across different games.

Our results are based on the assumption that individuals have an equal likelihood of finding themselves in a high-or low-power position. This assumption reflects the empirical observation that variations in people’s perceptions of power are mainly due to changes in situations, rather than their stable traits (21). In addition, we used variations of the simultaneous donation game as a framework to study the emergence of reciprocal cooperation. Including in our analysis both PD and SD games and varying the continuation probability allow us to explore a space of the four “archetypal” games most frequently studied in the literature on social dilemmas (54,57–59); yet, this is but a fraction of the 8-dimensional parameter space of all asymmetric, dyadic games. Further studies are needed to determine to what extent our conclusions can be generalized across all possible games.

Another important question concerns the exploration of alternative strategies of reciprocal cooperation. We only focused on TFT and AllD: this is because under the assumptions of our model (no noise and no costs associated with TFT compared to other strategies), if TFT cannot outcompete AllD, no other cooperative strategy can (7). While TFT may pave the way for cooperation to evolve, it is not necessarily the best strategy to maintain cooperation once it has been established (11,12). In fact, TFT can be invaded by AllC through drift, making it possible for AllD to eventually take over the population (11); on average, however, these evolutionary cycles tend to favour TFT (72). As the main focus of this work is to establish the necessary conditions for cooperation to emerge, and TFT invading AllD is the minimum requirement for cooperation to evolve (7), we did not consider other strategies than TFT and AllD. Future investigations on cooperation in asymmetric games could explore a broader space of strategies, such as zero-determinant strategies (73,74), win-stay lose-shift (11,12), or forgiving TFT(75), and study the impact of alternative update rules (76).

It is often said that “power corrupts”, expressing the common wisdom that individuals in high-power positions can succumb to the temptation to exploit their subordinates (77). Our results indicate that, when power is operationalized as the ability to provide higher rewards, power asymmetry makes it harder for reciprocal cooperation to evolve: as low-power individuals cannot provide enough benefits to their social partners to even out the costs of cooperation, high-power individuals have a stronger incentive to defect. However, conditioning one’s decision to cooperate on the level of power is not always an optimal strategy. In the iterated PD, it is more beneficial to adopt a power-independent strategy of either reciprocal cooperation (TFT) or defection (AllD), regardless of power status (Figures 2 and 3). In the iterated SD, on the other hand, power offers the means to maximize collective payoffs through anti-coordination: low-power individuals benefit from “carrying” the cost of cooperation, as they will receive the same treatment when they find themselves in a high-power position (Figures 4 and 5). Yet, regardless of whether the optimal strategy is conditional or not on the level of power, higher levels of asymmetry hinder the emergence of cooperation (Figure 1) and thus lowers cooperation rates (Figure 6). Power asymmetry undermines reciprocal cooperation (Figures 1 and 6), and interventions to enhance cooperative behaviour should aim at promoting a more egalitarian profitability of mutual cooperation.

## Methods

### Operationalization of power

Interdependence theory provides the means of quantifying the exact amount of power asymmetry of an interaction described by a given payoff matrix (46,47). The variance in outcomes of an interaction can be divided in three distinct components: actor control (AC), partner control (PC), and joint control (JC). These values indicate to what extent the variance in outcomes is determined, respectively, by the player’s choices, by their partner’s choices, or by the importance of coordination (that is, the difference between making the same choice as one’s social partner or doing the opposite of what they do).

Let 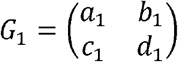 be the payoff matrix for player 1. Then 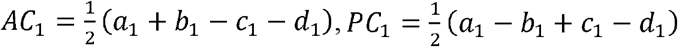 and 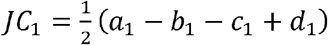 These quantities can be used to calculate the degree of mutual dependence (MD) of an interaction, which quantifies to what extent player 1’s payoff depends on player 2’s choices:

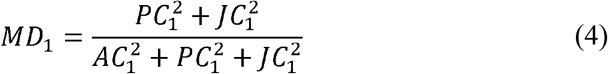

Mutual dependence varies between (no dependence) and (total dependence). In asymmetric interaction, this quantity can differ between the two players; for example, if *MD*_1_> *MD*_2_, player 2’s choices affect player 1’s outcome to a greater extent than player 1’s choices affect player 2’s outcome. The degree of asymmetric dependence, the difference in mutual dependence between the two players (*MD*_1_–*MD*_2_), indicates the asymmetric ability to control another person’s outcomes. As shown in Figure S1, this quantity increases monotonically with power asymmetry (*α*) as we defined it in the context of the asymmetric donation game.

### Evolutionary stability of TFT

Let *W* be the continuation probability (i.e., the likelihood of repeated encounters between the same individuals). In the PD, when only two strategies, TFT and AllD, are considered, and the interaction is characterized by power asymmetry *α*, the result is a bimatrix game with the following payoff structure:

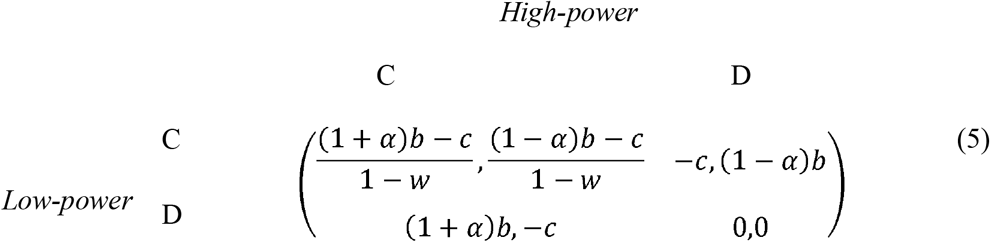

The necessary conditions for TFT to be evolutionarily stable for both players are 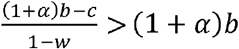 and 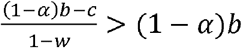, This leads to 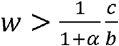 and 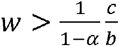 As *α >* 0 the first condition is always satisfied if the second is. Using a similar reasoning, it is possible to obtain the same conditions for the iterated SD (as the reward for mutual cooperation and the temptation to defect are the same in the two games, the conditions are the same). Here we focus on TFT and AllD alone as the evolutionary stability of TFT against AllD is a necessary condition for the evolution of reciprocal altruism (7).

### Evolutionary stability of strategies conditional on power

We consider an infinite population where individuals can randomly find themselves in a high-or low-power position, and interact with random members of the population. Therefore, every individual engages in symmetric interactions with a frequency of 0.5, in interactions with a lower-power individual with a frequency of 0.25, and in interactions with a higher power individual with a frequency of 0.25. The total payoff is averaged over a large number of interactions. We assume the population to be well-mixed. In each interaction, depending on the level of power asymmetry, an individual can choose whether to adopt reciprocal cooperation (TFT) or always defect (AllD). This gives rise to eight possible strategies: two “power-independent” strategies, where individuals always play TFT or AllD in every situation regardless power differences, and six strategies contingent on power differences, where individuals choose whether to play TFT or AllD depending on the specific type of interaction (symmetric or with a higher/lower-power individual). We do not consider strategies that distinguish between low-power and high-power symmetric interactions as we are mainly interested in studying the stability of strategies that depend on power asymmetries. Each of the eight strategies can then be identified as a triplet (*X,Y,Z*) where an individual plays *X* in symmetric interactions, *Y* when in low power and interacting with a higher-power individual, and *Z* when in high power and interacting with lower-power individual. From simplicity, we use the shorthand “C” to refer to reciprocal cooperation (TFT), and “D” as a shorthand for (*C,C,D*). For example, the strategy () plays TFT in symmetric interactions and when interacting with higher-power individuals, and AllD when interacting with individuals with a lower power status. Assuming that individuals have the same probability of being in a low or high-power state, and this state can eventually change between each interaction, the total payoff of an individual adopting strategy *i* is 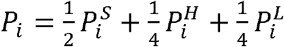, where 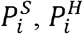 and 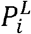 are, respectively, the average payoff of strategy in symmetric, high-power, and low-power interactions, which will in turn depend on the frequencies of other strategies in the population.

Let P_*ji*_ be the average payoff of a rare mutant playing strategy *j* in a population where strategy *i* is fixed. Strategy *i* is evolutionarily stable if, and only if, P_*ii*_ *>* P_*ij*_for every other strategy *j*, or if P_*ii*_= P_*ij*_ and P_*ii*_ *>* P_*ij*_ (66). We evaluated numerically which strategies are evolutionarily stable for given values of α and *w*.

### Small-mutation approximation

We also analyse the evolutionary dynamics of the strategies defined above in a finite population using a small mutation approximation, i.e., assuming that the time between the emergence of a new mutant playing a different strategy is much greater than the time it takes for a mutant to go extinct or spread to fixation (67). We model the evolutionary dynamics of the population as a Moran process, where at each time step an individual is randomly selected proportionally to their fitness to replace another, randomly selected individual (78,79). We define the fitness of an individual adopting strategy *i* as 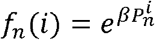, where *β* denotes the strength of selection and 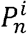 denotes the payoff of an individual playing strategy *i* is selected for reproduction is *x*(*i*)*f*_*n*_(*i*)/ Σ _*k*_ *x* (*k*) *f*_*n*_(*k*), where the denominator is the average fitness of the population. The offspring of the individual selected for reproduction replaces another individual, selected at random independently of fitness; that is, the probability that an individual adopting strategy *j* is replaced is *x* (*j*)/*N*.

We model the evolution of this population as a discrete time Markov chain with transition matrix *T* whose elements T_*ij*_indicate the transition probability from state *j*(where strategy *j*is fixed in the population) to state *i* (where strategy *i* is fixed) The transition probability from *j* to *i* is given by the product between population size (*N*), mutation rate (*U*) and the rate of evolution *P*_*ij*_ i.e., the probability of fixation of a single mutant adopting strategy ***I*** in a population of *j* (80). As the stationary distribution of a Markov chain does not change if all non-diagonal entries of the transition matrix are multiplied by the same factor, we can omit the terms *N* and *U* and write the transition coefficients as:

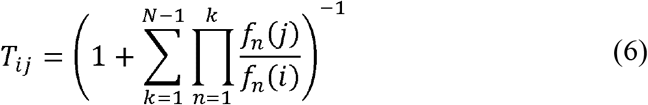

As there are no absorbing states, this matrix represents an ergodic Markov chain with a unique stationary distribution, which can be calculated as the left eigenvector associated with the eigenvalue λ =1 of the transition matrix.

### Exact stationary distribution

In order to evaluate whether the small-mutation approximation is a reasonable assumption to describe the evolution of such population, we consider the simplified case of only three strategies, for which the stationary distribution can be calculated exactly. For the PD, we considered the two power-independent strategies and one of the evolutionarily stable strategies conditional on power,(*C,D,D*) For the SD, we considered the two power-independent strategies and the only evolutionarily stable strategy conditional on power, (*C,C,D*) These strategies were selected as they can be evolutionary stable for certain values of *w* and α in an infinite population (Figure 2 and 4) and outcompete other conditional strategies in a finite population (Figure 3). The population evolves according to a Moran process, as described in the previous sections, with the only difference that, every time an individual reproduces, it mutates to a different strategy with probability *U*. At each time step, an individual adopting strategy *i* is selected with probability *x*(*i*)*f*_*n*_(*i*)/ Σ_*k*_*x* (*k*) *f*_*n*_(*k*), probability (1-*U*) their offspring adopts strategy *i* With probability *U*, the offspring adopts a different strategy (the offspring has an equal probability of adopting any other strategy). As previously, the offspring replaces a randomly chosen individual, independently of their fitness.

Let (*k,p,q*) denote a state with *k* individuals adopting (*C,C,C*),*p* individuals adopting(*D,D,D*) and *q* individuals adopting (*C,C,D*) and let *x=k/N,y=p/N*, and *z=q/N* the frequencies of these strategies in the population. The transition probability from (*k,p,q*) to (*k+*1,*p-*1,*q*) is:

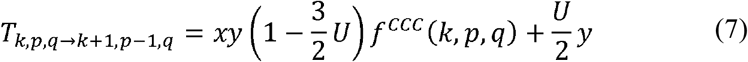

Where *f*^*ccc*^ designs the relative fitness of (*C,C,C*) in state(*k,p,q*). Analogous formulae describe the transitions from (*k,p,q*) to five other possible adjacent states. The transition probabilities are calculated numerically, and the transition matrix is then used to calculate the stationary distribution. As shown in Figures S4 and S5, up to a mutation rate of *U=*0.001 the system spends a negligible time in the intermediate states, the pure states (where one strategy has reached fixation in the population) being the ones with the highest frequency (Figures S4 and S5). This supports our choice of adopting the small-mutation approximation described in the previous paragraph.

## Acknowledgments

This was supported by the European Research Council (ERC) under the European Union’s Horizon 2020 research and innovation programme (Grant agreement No. 864519), awarded to Daniel Balliet.

