## Supplementary Information for "Power asymmetry destabilizes reciprocal cooperation in social dilemmas"

*SI Appendix A – Transition probabilities in a 3-strategy Moran process with mutation*

Let $(k,p,q)$ denote a state with $k$ individuals adopting $(C,C,C)$, $p$ individuals adopting $(D,D,D)$ and $q$ individuals adopting $(C,C,D)$. Let $x=k/N$, $y=p/N$, and $z=q/N$ be the frequencies of these strategies in the population, and $f^{CCC}$, $f^{DDD}$, and $f^{CCD}$ their *relative* fitness in state $(k,p,q)$, i.e., $f^{X}=F^{X}/\bar{f}$ (where $F^{X}$is the absolute fitness of strategy $X$ and $\bar{f}=xF^{CCC}+yF^{DDD}+zF^{CCD}$ is the average fitness in the population). Therefore, ${xf}^{CCC}+{yf}^{DDD}+{zf}^{CCD}=1$.

Assuming that strategies evolve according to a Moran Process, the transition probability from $(k,p,q)$ to $(k+1,p-1,q)$ is the probability that a $(C,C,C)$ individual replaces a $(D,D,D)$ individual without mutating, or that a $(D,D,D)$ individual is replaced by either a $(D,D,D)$ or a $(C,C,D)$ individual which mutates to $(C,C,C)$. This can be written as:

$$T_{k,p,q\to k+1,p-1,q}=\frac{p}{N}\left[ \left( 1-U \right)\frac{k}{N}f^{CCC}+\frac{U}{2}\frac{p}{N}f^{DDD}+\frac{U}{2}\frac{q}{N}f^{CCD} \right]=y\left[ \left( 1-U \right)xf^{CCC}+\frac{U}{2}yf^{DDD}+\frac{U}{2}zf^{CCD} \right]=y\left[ xf^{CCC}-Uxf^{CCC}+\frac{U}{2}\left( yf^{DDD}+zf^{CCD} \right) \right]=y\left[ xf^{CCC}-Uxf^{CCC}+\frac{U}{2}\left( 1-xf^{CCC} \right) \right]=y\left( xf^{CCC}-Uxf^{CCC}+\frac{U}{2}-\frac{U}{2}xf^{CCC} \right)=xy\left( 1-\frac{3}{2}U \right)f^{CCC}\left( k,p,q \right)+\frac{U}{2}y$$

Transition probabilities s to all other adjacent states can be calculated in a similar way.

**Supplementary Figures**

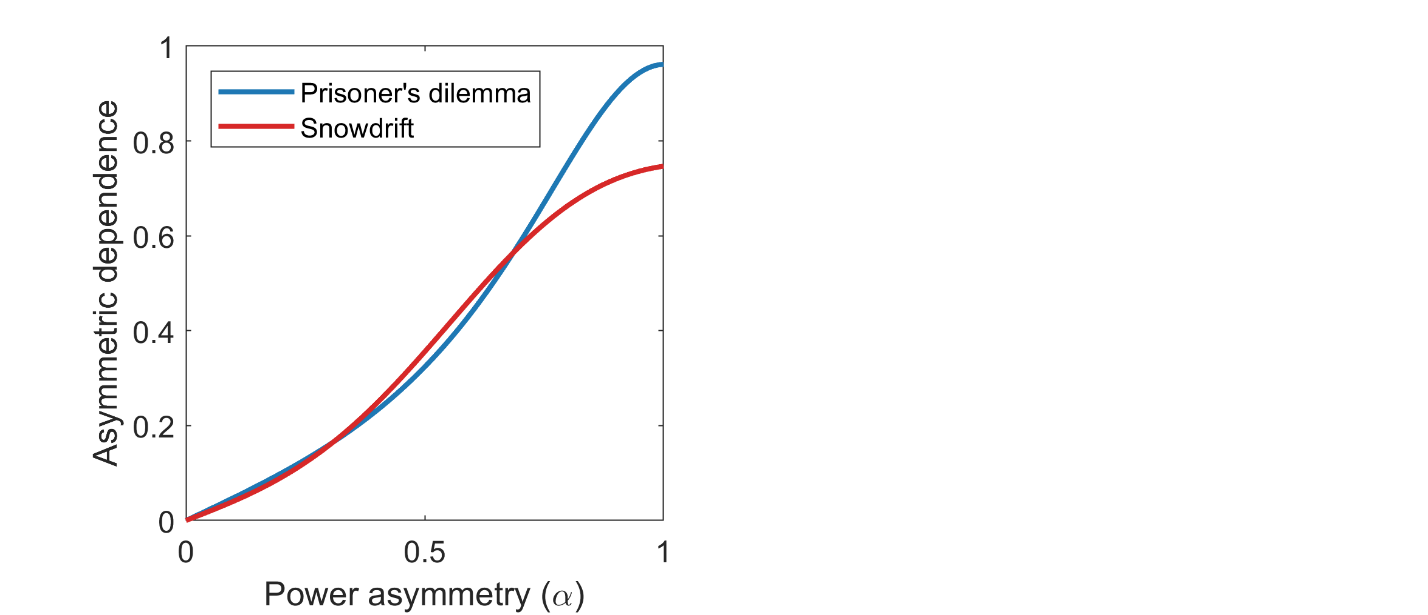

**Fig. S1.**

Relationship between power asymmetry ($\alpha)$ the degree of asymmetric dependence, as defined in interdependence theory (Balliet et al., 2017; Kelley et al., 2003), in the PD and SD. Parameters: $b=1$, $c=4$, $w=0$.

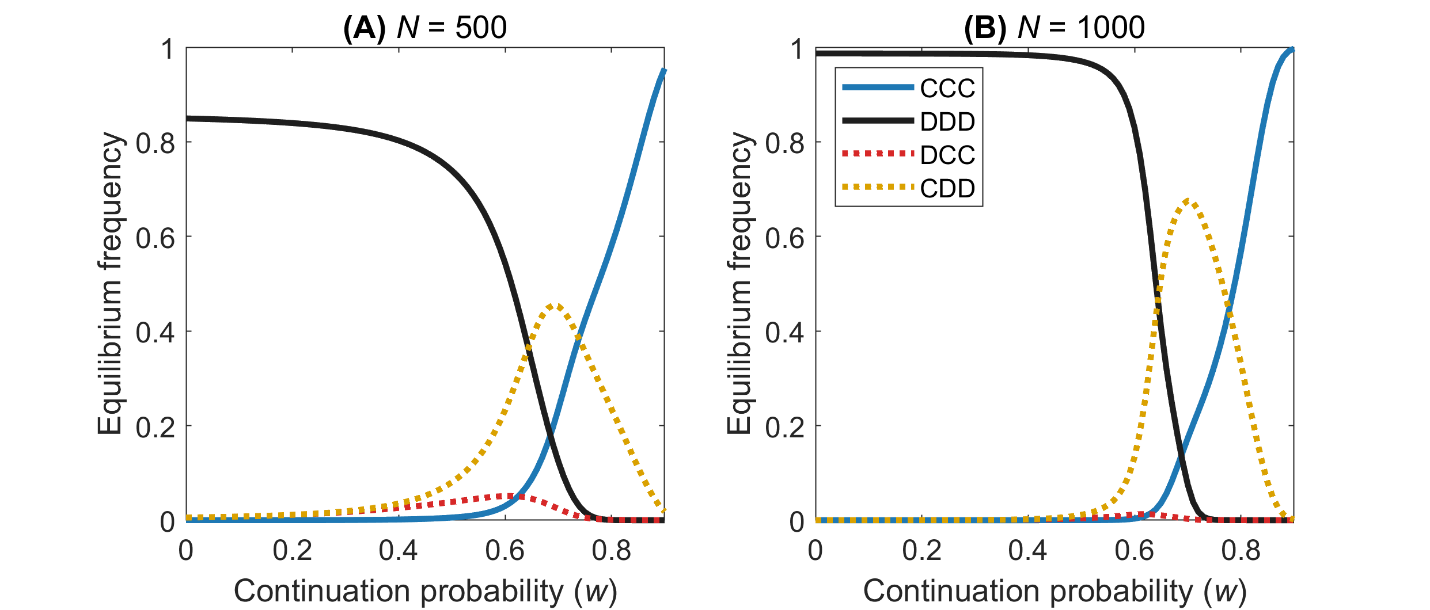

**Fig. S2.** **Equilibrium frequency of strategies in the iterated PD.** Equilibrium frequency of various strategies in a finite population ($N=500$ (**A**) and $N=1000$ (**B**)) playing the iterated PD, under a small-mutation approximation, as a function of the continuation probability ($w$). Continuous lines indicate power-independent strategies; dotted lines indicate strategies that are conditional on power differences. Of the 8 strategies studied, only the four ESSs are shown. Other parameters: $\alpha=0.5$, $b=1$, $c=0.4$, $\beta=0.05$.

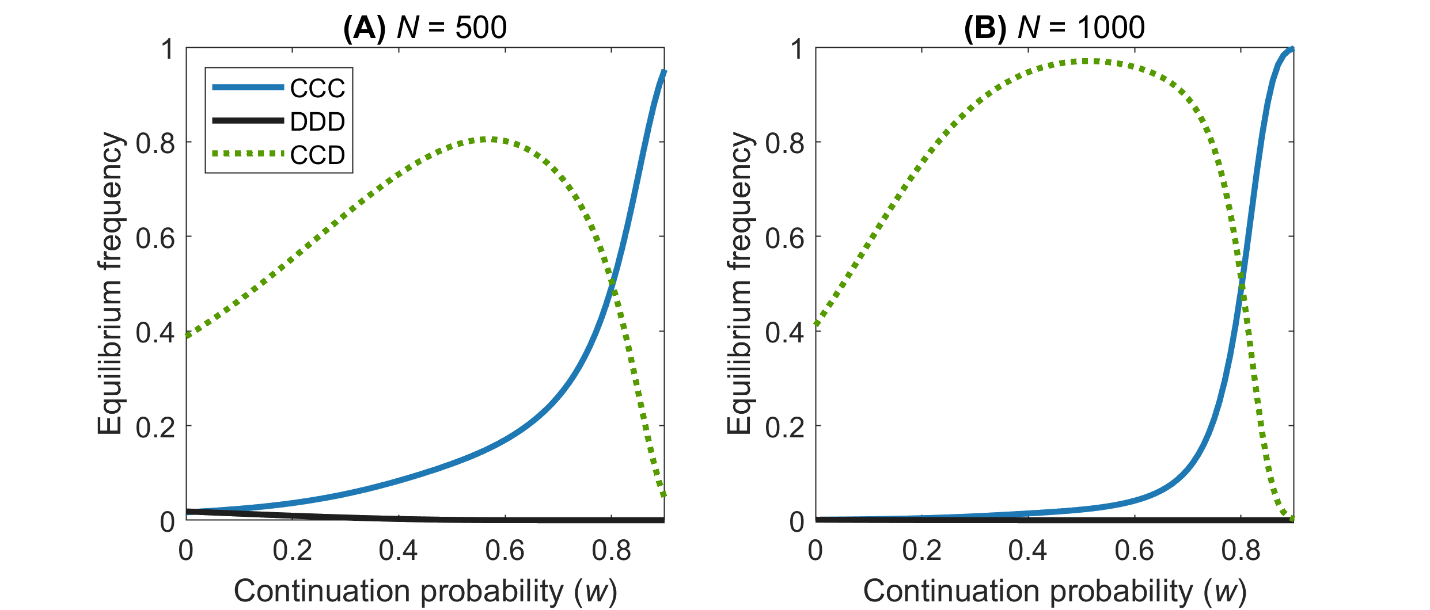
**Fig. S3.** **Equilibrium frequency of strategies in the iterated SD.** Equilibrium frequency of various strategies in a finite population ($N=500$ (**A**) and $N=1000$ (**B**)) playing the iterated SD, under a small-mutation approximation, as a function of the continuation probability ($w$). Continuous lines indicate power-independent strategies; the dotted line indicates ($C,D,D$), a strategy that is conditional on power differences. Of the 8 strategies studied, only the two ESSs and AllD are shown. Other parameters: $\alpha=0.5$, $b=1$, $c=0.4$, $N=100$, $\beta=0.05$.

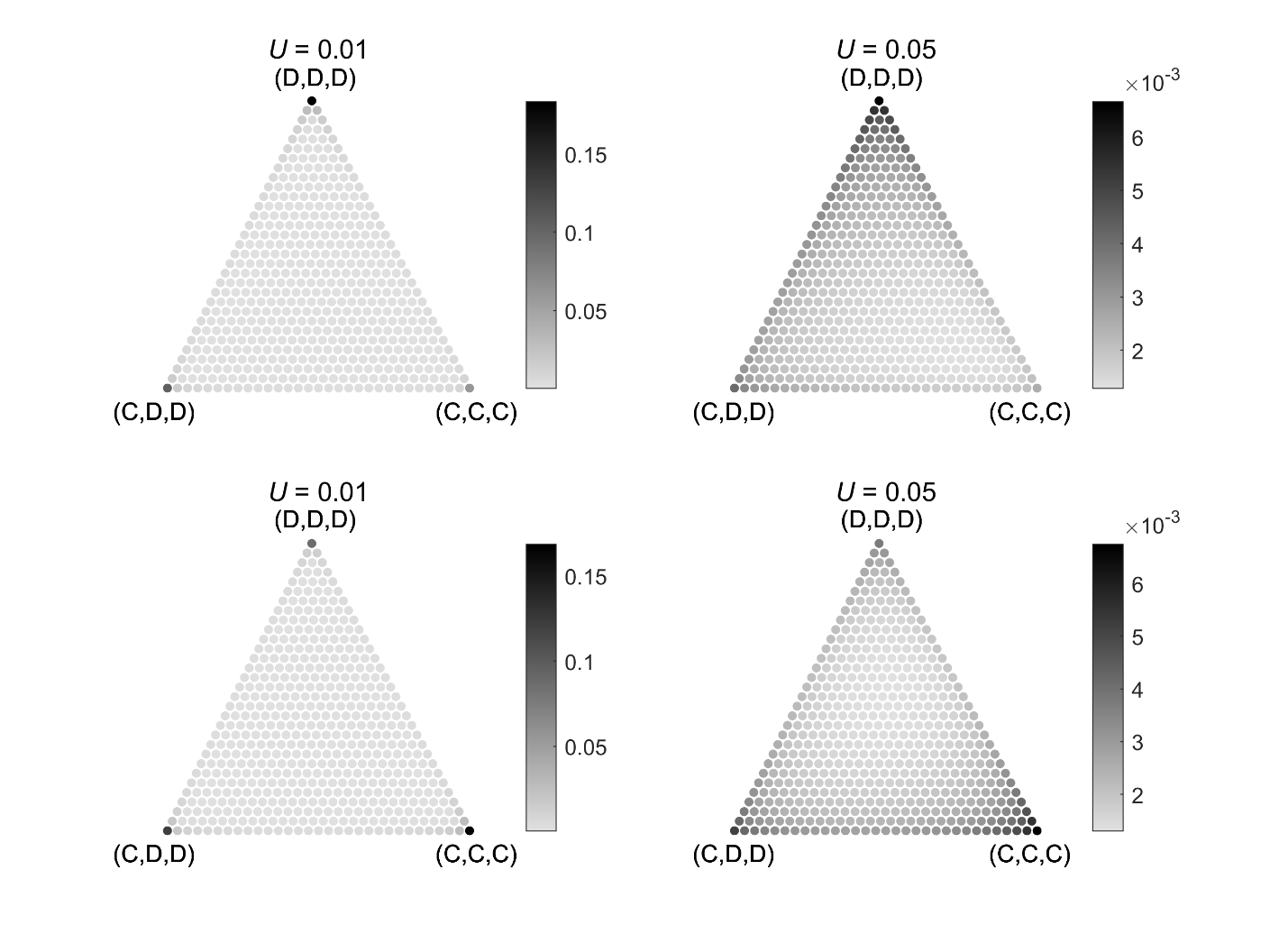

**Fig. S4.**

Stationary distribution over strategies in a finite population composed of $(C,C,C)$, $(D,D,D$), and ($C,D,D$) and playing the iterated PD. The stationary distributions are calculated as the limiting distribution of a discrete-time Markov chain (which, as long as U>0, is irreducible, thereby has a unique stationary distribution). For a mutation rate $U=0.01$ (left two graphs), the population spends most of the time in pure states, validating the use of a small-mutation approximation. When the continuation probability is low ($w=0.1$; upper two graphs), defecting regardless the level of power is the optimal strategy. When the probability of repeated encounters is high ($w=0.9$), reciprocal cooperation is the optimal strategy, regardless the level of power of an interaction. The fixation probability of $(D,C,C)$ in a population of $(C,C,C)$, $(D,D,D$), and ($D,C,C$) is the same as $(C,D,D)$ and therefore not shown as a separate figure. Other parameters: $\alpha=0.5$, $b=1$, $c=0.4$, $N=30$, $\beta=0.05$.

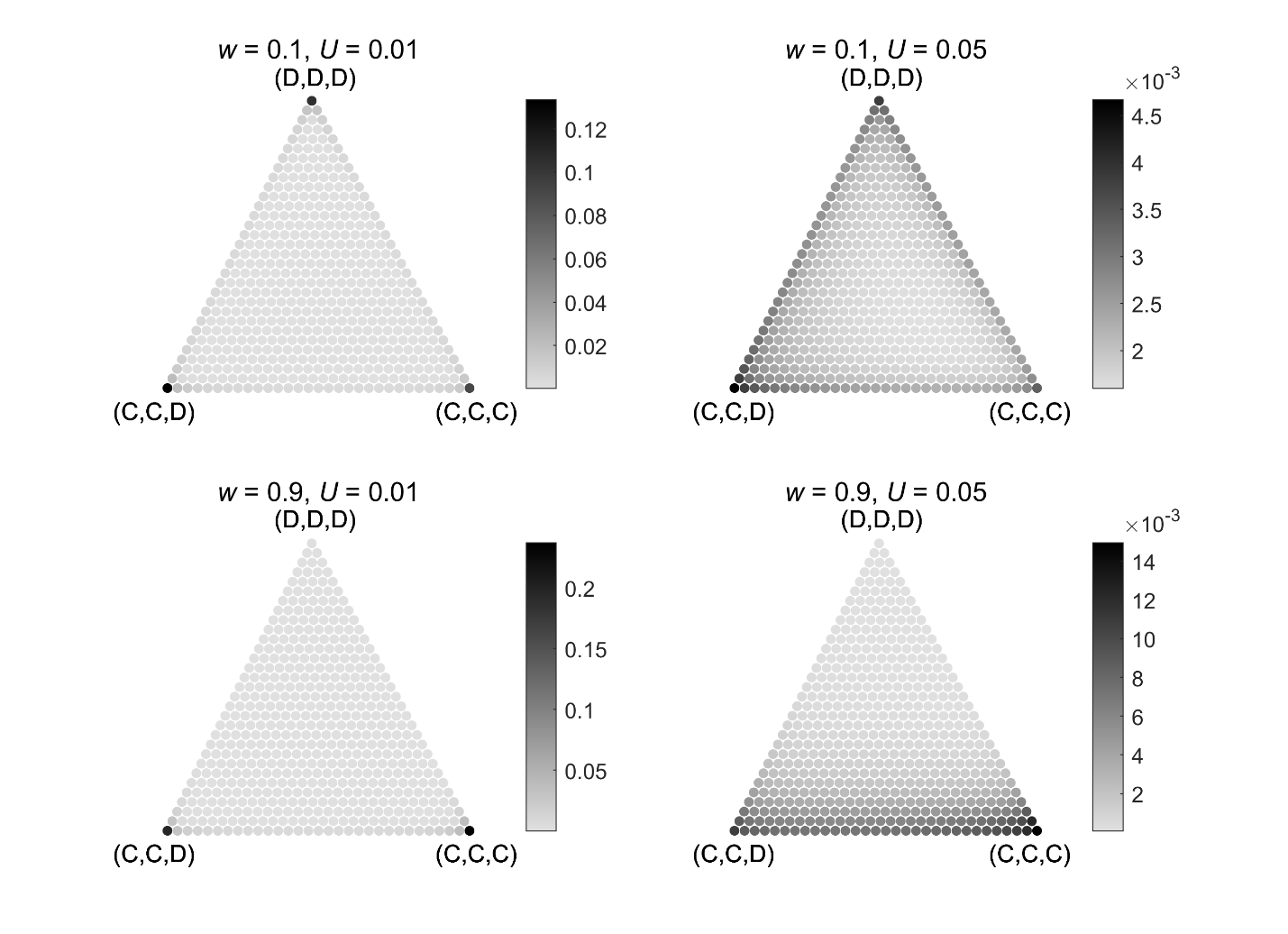

**Fig. S5.**

Stationary distribution over strategies in a finite population composed of $(C,C,C)$, $(D,D,D$), and ($C,C,D$) and playing the asymmetric, iterated SD. The stationary distributions are calculated as the limiting distribution of a discrete-time Markov chain. For a mutation rate $U=0.01$ (left two graphs), the population spends most of the time in pure states, validating the use of a small-mutation approximation. When the continuation probability is low ($w=0.1$; upper two graphs), ($C,C,D$) is the optimal strategy. When the probability of repeated encounters is high ($w=0.9$), reciprocal cooperation is the optimal strategy, regardless the level of power of an interaction. Other parameters: $\alpha=0.5$, $b=1$, $c=0.4$, $N=30$, $\beta=0.05$.
